# TRPS1 modulates chromatin accessibility to regulate estrogen receptor (ER) binding and ER target gene expression in luminal breast cancer cells

**DOI:** 10.1101/2023.07.03.547524

**Authors:** Thomas G. Scott, Kizhakke Mattada Sathyan, Daniel Gioeli, Michael J. Guertin

## Abstract

Breast cancer is the most frequently diagnosed cancer in women. The most common subtype is luminal breast cancer, which is typically driven by the estrogen receptor *α* (ER), a transcription factor (TF) that activates many genes required for proliferation. Multiple effective therapies target this pathway, but individuals often develop resistance. Thus, there is a need to identify additional targets that regulate ER activity and contribute to breast tumor progression. TRPS1 is a repressive GATA-family TF that is overexpressed in breast tumors. Common genetic variants in the TRPS1 locus are associated with breast cancer risk, and luminal breast cancer cell lines are particularly sensitive to TRPS1 knockout. However, we do not know how TRPS1 regulates target genes to mediate these breast cancer patient and cellular outcomes. We introduced an inducible degron tag into the native TRPS1 locus within a luminal breast cancer cell line to identify the direct targets of TRPS1 and determine how TRPS1 mechanistically regulates gene expression. We acutely deplete over eighty percent of TRPS1 from chromatin within 30 minutes of inducing degradation. We find that TRPS1 regulates transcription of hundreds of genes, including those related to estrogen signaling. TRPS1 directly regulates chromatin structure, which causes ER to redistribute in the genome. ER redistribution leads to both repression and activation of dozens of ER target genes. Downstream from these primary effects, TRPS1 depletion represses cell cycle-related gene sets and reduces cell doubling rate. Finally, we show that high TRPS1 activity, calculated using a gene expression signature defined by primary TRPS1-regulated genes, is associated with worse breast cancer patient prognosis. Taken together, these data suggest a model in which TRPS1 modulates the activity of other TFs, both activating and repressing transcription of genes related to cancer cell fitness.

## Introduction

Breast cancer is the most frequently diagnosed cancer in women, with an estimated lifetime risk of about 1 in 8 for women in the United States (Giaquinto et al. 2022). Far from a monolithic disease, breast tumors can be classified into subtypes based on gene expression, histology, and immunohistochemistry (Cheang et al. 2015; Perou et al. 2000). The most common subtype is luminal breast cancer, which is typically estrogen receptor *α* (ER)-positive (Harbeck et al. 2019).

High lifetime exposure to endogenous estrogen is a strong risk factor for breast cancer incidence (Henderson et al. 1982). Estrogen is a potent hormone that binds to ER, a ligand-activated transcription factor (TF), which then ho-modimerizes, binds to reverse palindromic pairs of AG-GTCA motifs on DNA, and recruits cofactors to activate hundreds of genes that promote cell growth and proliferation (Hah et al. 2011; Hall et al. 2001; Kumar and Chambon 1988). In additional to surgery, radiation, and traditional chemotherapy, endocrine therapies that inhibit endogenous estrogen production or binding to ER provide a significant survival benefit to luminal breast cancer patients (Francis et al. 2018; Gnant et al. 2015; Harbeck et al. 2019). However, patients with advanced disease frequently develop resistance to these therapies, though many endocrine therapy-resistant luminal tumors still remain dependent on ER activity (Dodwell et al. 2006). Thus, there is a need to identify additional factors that regulate ER activity and contribute to breast tumor progression.

TRPS1 is a member of the GATA-family of TFs that bind to (A/T)GATA(A/G) motifs on DNA (Ko and Engel 1993). In contrast to the other six members of the GATA family that activate transcription, TRPS1 directly represses transcription of target genes via its unique Ikaros-like zinc fingers (Malik et al. 2001). TRPS1 has been shown to interact with multiple corepressors and lysine deacetylases, including members of the NuRD and coREST complexes, to regulate transcription (Cornelissen et al. 2020; Elster et al. 2018; Serandour et al. 2018; Wang et al. 2018a,b).

TRPS1 was first described as the gene mutated in cases of tricho-rhino-phalangeal syndrome, an autosomal dominant disorder characterized by developmental abnormalities of the hair, nose, and fingers (Momeni et al. 2000). TRPS1 is crucial for the proper development of several tissues, including hair, bone, and kidney (Gai et al. 2009; Malik et al. 2002). As with many developmentally important genes coopted during the process of cancer initiation and progression, TRPS1 is commonly over-expressed in breast tumors, both relative to normal tissue and relative to other tumor types (Ai et al. 2021; Lin et al. 2017).

The transcriptional program that TRPS1 regulates in breast cancer is not fully understood. Knockdown of TRPS1 in various breast cancer cell lines has been shown to increase markers of epithelial to mesenchymal transition and genome instability (Hu et al. 2018; Huang et al. 2016; Stinson et al. 2011; Yang et al. 2021). Additionally, TRPS1 binding sites on chromatin overlap with those of YAP and ER, though this is coupled with a genome-wide activation of YAP target genes but repression of ER target genes (Elster et al. 2018; Serandour et al. 2018).

A key feature of these previous studies is the use of extended knockdown kinetics with traditional RNA interference methods. Days after knockdown, the resultant effects represent not only the primary TRPS1-responsive genes but also secondary and compensatory effects. As such, we do not know which genes TRPS1 directly regulates and whether these genes are important for breast cancer cell growth and proliferation.

In this study, we set out to directly assay the primary effects of TRPS1 on chromatin accessibility, ER binding, and transcription in luminal breast cancer cells. To do so, we acutely depleted TRPS1 using an inducible degron tag inserted into the endogenous TRPS1 locus. By performing sensitive genome-wide assays minutes to hours after TRPS1 depletion, we demonstrated that TRPS1 changes chromatin structure, which allows ER to redistribute in the genome. Through this redistribution, TRPS1 both directly represses and indirectly activates dozens of ER target genes at baseline. Furthermore, we defined a signature of primary TRPS1-regulated genes that predicts breast cancer patient prognosis.

## Results

### TRPS1 is associated with breast cancer incidence and promotes breast cancer cell number accumulation

A recent genome-wide association study (GWAS) identified 32 novel single nucleotide polymorphisms (SNPs) associated with breast cancer susceptibility (Zhang et al. 2020). When we queried the NHGRI EBI GWAS Catalog to find published associations with genetic variants within the TRPS1 genomic locus, we found one of the lead SNPs from this study (Sollis et al. 2023). Furthermore, in the authors’ subtype-specific analysis of the results, the association with this variant was strongest for luminal breast cancers relative to other subtypes (Zhang et al. 2020).

We used LocusZoom to plot the data within this locus (Figure 1A). A plot of these data indicates that two sets of SNPs have low linkage disequilibrium with one another, indicating that they are inherited independently and each confer risk. One set of variants is within an intronic region of the TRPS1 gene, and one is about 400 kilobases upstream from the transcription start site (TSS) of the TRPS1 gene.

**Fig. 1.**
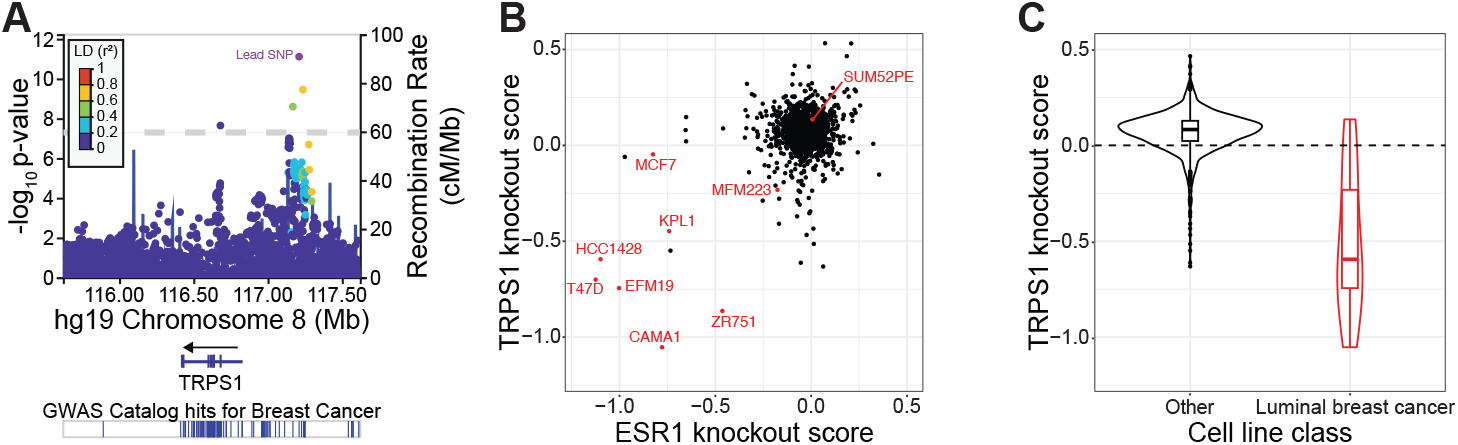
TRPS1 is associated with breast cancer incidence and promotes breast cancer cell fitness. A) LocusZoom plot of the TRPS1 genomic locus depicting the location and significance of SNPs associated with breast cancer susceptibility. TRPS1 is the closest gene to two sets of genetic variants in low linkage disequilibrium with one another. Data from (Zhang et al. 2020), generated with summary statistics downloaded from the NHGRI-EBI GWAS Catalog, using LocusZoom (Boughton et al. 2021; Sollis et al. 2023). B) Scatter plot of TRPS1 and ESR1 knockout scores for each gene tested. Scores are normalized such that knockout of a gene with a score of 0 has no effect on cell number, and knockout of a gene with a score of -1 has an effect equal to that of knocking out one of a set of universally essential genes. Luminal breast cancer cell lines are colored in red. The data are from the Cancer Dependency Map project (Meyers et al. 2017). C) Violin and box and whisker plots of TRPS1 knockout scores from (B) for luminal breast cancer cell lines versus all other cell lines. Wilcoxon rank sum test p-value of 3.2*10^−5^.

There are often many genes in close proximity to the lead SNP in a GWAS, making it difficult to predict which gene mediates the effect on the associated phenotype. However, in this case the nearest gene is almost a megabase upstream from TRPS1, suggesting that TRPS1 itself contributes to the breast cancer susceptibility associated with one or both of these sets of genetic variants.

Based on this result, we hypothesized that perturbation of TRPS1 in luminal breast cancer cell lines would affect cell fitness. Using data from the Cancer Dependency Map project, we found that sensitivity to TRPS1 knockout correlated with sensitivity to knockout of ESR1, the gene encoding ER (Figure 1B) (Meyers et al. 2017). Furthermore, while TRPS1 knockout led to an increase in cell number for most cancer cell lines, luminal breast cancer cell lines were significantly enriched for TRPS1 dependency (Figure 1C).

Taken together, these data indicate that TRPS1 influences breast cancer incidence and is required for maximal breast cancer cell fitness. Next we sought to determine how TRPS1 regulates its target genes to mediate these breast cancer patient and cellular outcomes.

### TRPS1 that is endogenously degron-tagged is rapidly degraded in T47D cells

To rapidly deplete TRPS1 and isolate primary TRPS1-regulated genes, we employed the dTAG system for targeted protein degradation (Nabet et al. 2020, 2018). We inserted an inducible degron tag into the endogenous TRPS1 locus in the luminal breast cancer cell line T47D. We generated three independent clones that express the tagged TRPS1 and can be degraded by the addition of the small molecules dTAG-13 and dTAG^V^-1 at 50nM each (dTAG) (Figure 2A). Of note, we depleted around 50% of TRPS1 from whole cell lysates in 10 minutes of treatment with dTAG, as determined by quantitative western blot, with less than 10% detected as soon as 20 minutes and as late as 48 hours after treatment (Figure 2B).

**Fig. 2.**
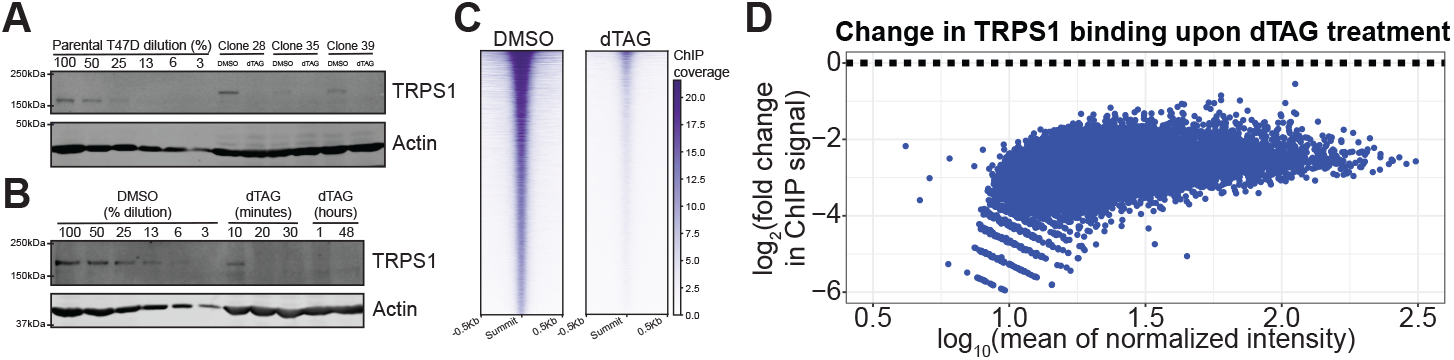
Endogenous tagging allows for rapidly inducible TRPS1 depletion in T47D cells. A) Quantitative Western blot with a serial dilution of the parental T47D cells followed by three independent dTAG-TRPS1 clones treated with DMSO or dTAG-13 and dTAG^V^-1 at 50nM each (dTAG) for 2 hours. Membranes were probed with anti-TRPS1 or anti-Actin antibodies. B) Quantitative Western blot with a serial dilution of dTAG-TRPS1 Clone 28 followed by a time course of treatment with dTAG. Membranes were probed as in (A). C) Heatmap of TRPS1 ChIP-seq peaks, in rows ranked by intensity, in cells treated with DMSO or dTAG for 30 minutes. D) MA plot of TRPS1 ChIP-seq peaks, with fold change values representing binding intensity in the dTAG condition relative to the DMSO condition. All points are colored blue to indicate they are significantly decreased at an FDR of 0.1.

To ensure this treatment depleted TRPS1 from chromatin, we performed chromatin immunoprecipitation with sequencing (ChIP-seq) using an anti-HA antibody to recognize the 2xHA tag within the degron tag. We observed a genome-wide decrease in TRPS1 binding intensity, with over 80 percent of TRPS1 depleted from chromatin after 30 minutes of dTAG treatment (Figure 2C,D).

### TRPS1 directly represses regulatory element activity

With this system in hand, we set out to test the effects of TRPS1 depletion on regulatory element (RE) (i.e., enhancer or promoter) activity. To capture the dynamics of chromatin accessibility after TRPS1 depletion, we conducted a time course analysis using the Assay for Transposase-Accessible Chromatin with sequencing (ATAC-seq). The time points included dTAG treatments at 30 minutes, 1 hour, 2 hours, 4 hours, and 24 hours, while a DMSO treatment served as the vehicle control, assigned as the zero minute time point.At the earliest time point after degradation, our best estimate of the primary effects of TRPS1 depletion, we identify many peaks that increased in intensity and fewer that decreased in intensity, at a false discovery rate (FDR) of 0.1. (Figure 3A). We hypothesized that the increased peaks result from loss of a direct TRPS1 repression of chromatin accessibility, with the decreased peaks an indirect effect of the redistribution of limiting cofactors.

**Fig. 3.**
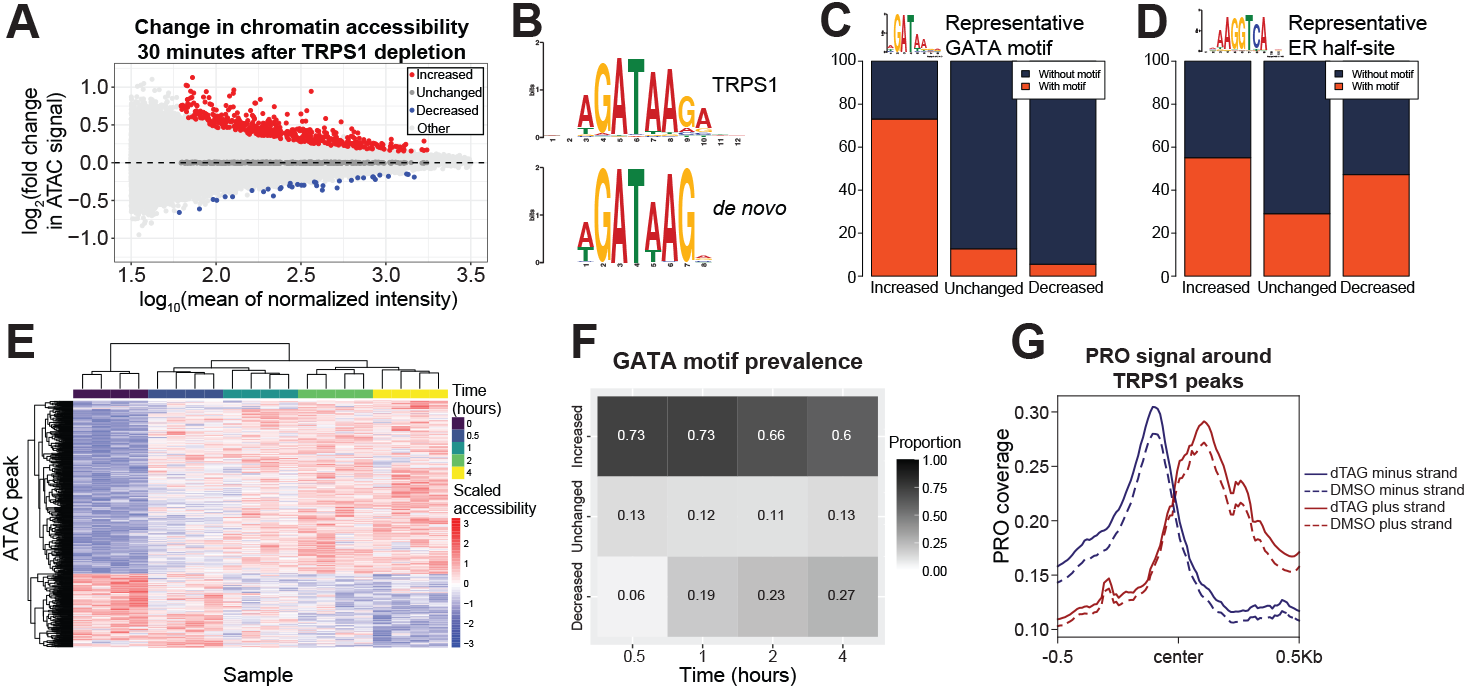
TRPS1 directly represses regulatory element activity. A) MA plot of ATAC-seq peaks, with fold change values representing accessibility in the 30 minute dTAG-13 and dTAG^V^-1 at 50nM each (dTAG) treatment condition relative to the DMSO condition. B) *De novo* motif identified in increased peaks from (A) (below), matched to the TRPS1 motif (above). C) Bar charts of prevalence of a representative GATA motif in increased, unchanged, and decreased peaks from (A). Chi-square test p-value < 2.2*10^−16^. D) Bar charts of prevalence of a representative ER half-site in increased, unchanged, and decreased peaks from (A). Chi-square test p-value 4.5*10^−15^. E) Heat map with hierarchical clustering of chromatin accessibility in ATAC-seq peaks that are significantly changed over the time course at an FDR of 0.1. F) Heat map of prevalence of the representative GATA motif in increased, unchanged, and decreased peaks, as in (C), for each time point relative to the DMSO condition. G) Density plot for composite PRO-seq signal across all TRPS1 ChIP-seq peaks, separated by strand and treatment condition.

To test this hypothesis, we performed *de novo* motif identification in the increased and decreased peaks. We identified a GATA motif in the increased peaks but not the decreased peaks (Figure 3B). To explicitly calculate the motif prevalence in each class of peaks, we found individual motif occurrences genome-wide and intersected the peaks with these motif instances. We found a significant enrichment of a representative GATA motif in the increased peaks over the unchanged and decreased peaks (Figure 3C).

To identify additional TFs that might be contributing to changes in chromatin accessibility, we analyzed motif enrichment in both increased peaks relative to unchanged peaks as well as decreased peaks relative to unchanged peaks (Tables S1, S2). To our surprise, we identified nuclear receptor motifs in both classes of peaks. To follow up this analysis, we again calculated motif prevalence in each class of peaks for a representative ER half-site. For this motif, we found a significant enrichment in both increased and decreased peaks relative to unchanged peaks (Figure 3D).

We predicted that the enrichment of the GATA motif within increased peaks would wane over time. To test this prediction, we turned to our time course data. We performed hierarchical clustering of ATAC-seq peaks that were significantly changed over the time course at an FDR of 0.1.

We found that the replicates for each time point clustered together and that the majority of dynamic peaks changed gradually over the time course, with additional clusters displaying different kinetics (Figure 3E). We called increased, unchanged, and decreased peaks as in Figure 3A for each time point relative to the control condition. We then calculated GATA motif prevalence in each set of peaks as in Figure 3C. As we expected, the GATA motif prevalence in the increased peaks decreased over time, but the majority of increased peaks at 4 hours still contained a GATA motif (Figure 3F). This is consistent with the primary effects of TRPS1 depletion driving a large proportion of the changes in chromatin accessibility even as late as 4 hours after TRPS1 depletion.

As an orthogonal readout of RE activity, we measured bidirectional transcription around TRPS1 binding sites using precision genomic run-on with sequencing (PRO-seq). Our libraries were of relatively high quality using several quality control metrics (Figure S1) (Scott et al. 2022). Using a window centered on each summit of TRPS1 ChIP-seq intensity, we observed an increase in bidirectional transcription 30 minutes after TRPS1 depletion (Figure 3G). Along with our accessibility data and motif analysis, these data indicate that TRPS1 directly represses RE activity.

### TRPS1 directly represses transcription of target genes

Downstream from the changes in RE activity, we measured changes in nascent transcription within genes with PRO-seq over the same time course of TRPS1 depletion as in the ATAC-seq experiment. We identified hundreds of dynamic genes over the time course and performed hierarchical clustering to classify the genes based on their expression ki-netics (Figure 4A, B). Over-representation analysis (ORA) of the activated genes identified several enriched Hallmark gene sets (Liberzon et al. 2015), most prominently cholesterol homeostasis genes (Figure 4C). ORA on the repressed genes revealed that the two estrogen response gene sets were the most significantly enriched of the Hallmark gene sets.

**Fig. 4.**
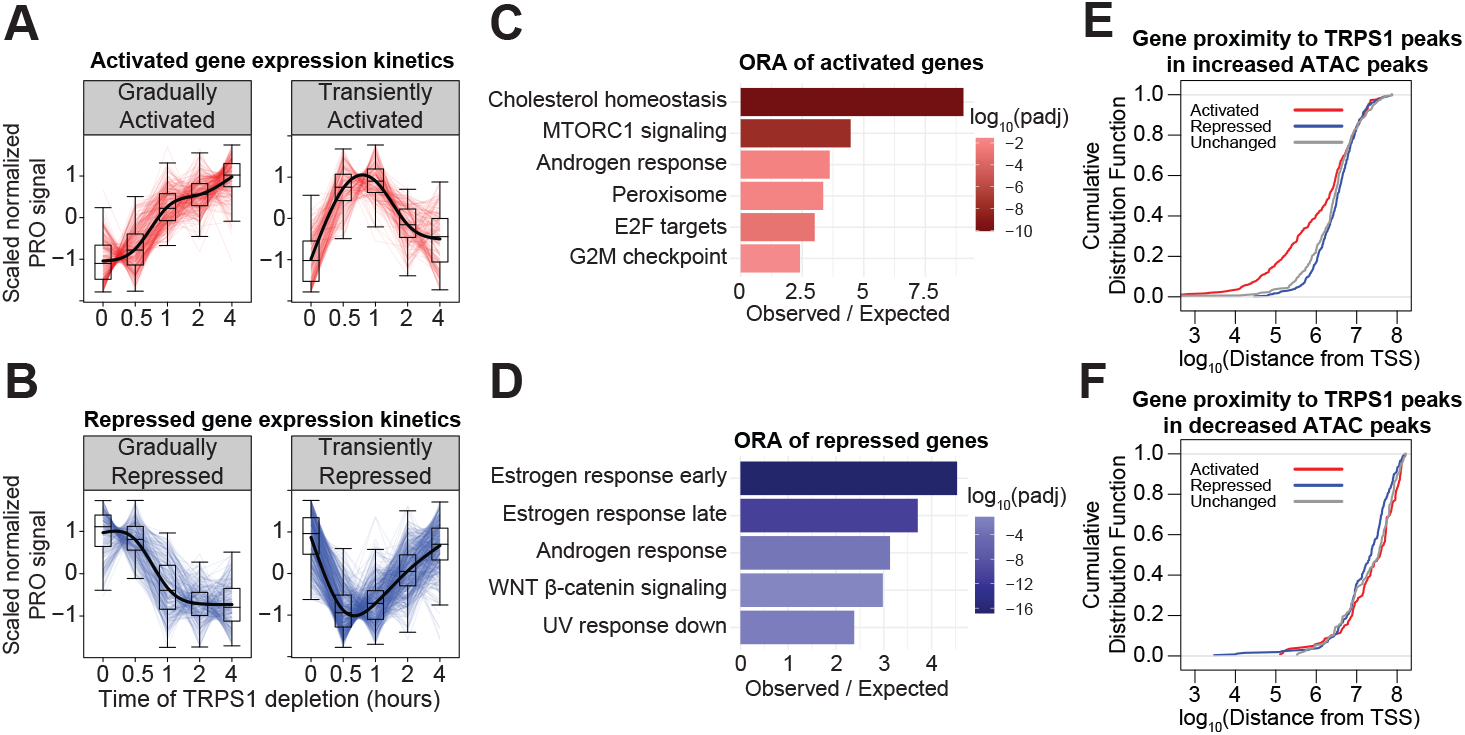
TRPS1 directly represses transcription of target genes. A) Kinetic traces of the two major clusters of activated genes over the time course. B) Kinetic traces of the two major clusters of repressed genes over the time course. C) Over-representation analysis of the activated genes from (A). D) Over-representation analysis of the repressed genes from (B). E) Cumulative distribution function plot of the distance from the TSS of each gene to the nearest TRPS1 ChIP-seq peak overlapping an increased ATAC-seq peak, by gene class. Kolmogorov–Smirnov test between activated and unchanged genes: p-value = 0.011. F) Cumulative distribution function plot of the distance from the TSS of each gene to the nearest TRPS1 ChIP-seq peak overlapping an decreased ATAC-seq peak, by gene class. KS test between repressed and unchanged genes: p-value > 0.1.

We hypothesized that TRPS1 regulates these gene sets by distinct mechanisms. Specifically, we predicted that TRPS1 In contrast to the genes activated by TRPS1 depletion, we predicted that the effects on the genes repressed by TRPS1 depletion are indirect and distal to TRPS1 binding. To test whether TRPS1 directly activates a subset of REs to activate transcription of proximal genes, we measured the distance from the TSS of each gene to the nearest TRPS1 ChIP-seq peak overlapping a decreased ATAC-seq peak. There are few examples of this class of RE, so the distances were much farther, and there was no significant enrichment of repressed genes proximal to these peaks (Figure 4F). These data suggest that, while TRPS1 positively and negatively regulates hundreds of primary response genes, TRPS1 only represses transcription of its direct target genes.

### TRPS1 redistributes ER binding to modulate ER target gene transcription

Based on the correlation between cancer cell line sensitivity to TRPS1 knockout and ER knockout (Figure 1B), the enrichment of an ER binding motif in both increased and decreased ATAC-seq peaks (Figure 3D), and the over-representation of estrogen response gene sets in the genes repressed by TRPS1 depletion (Figure 4D), we focused on ER target genes to explore the mechanism by which TRPS1 indirectly activates transcription. We first defined direct ER target genes using our previously-generated PRO-seq data from parental T47D cells that were hormone starved for three days and then acutely stimulated with estrogen or a DMSO vehicle control for 90 minutes (Figure S2) (Scott et al. 2022). We exclusively focused on estrogen-activated genes because ER directly activates these genes (Carroll et al. 2005; Guertin, MJ et al. 2014; Hah et al. 2011). Dozens of ER target genes were activated and repressed by acute TRPS1 depletion (Figure 5A).

**Fig. 5.**
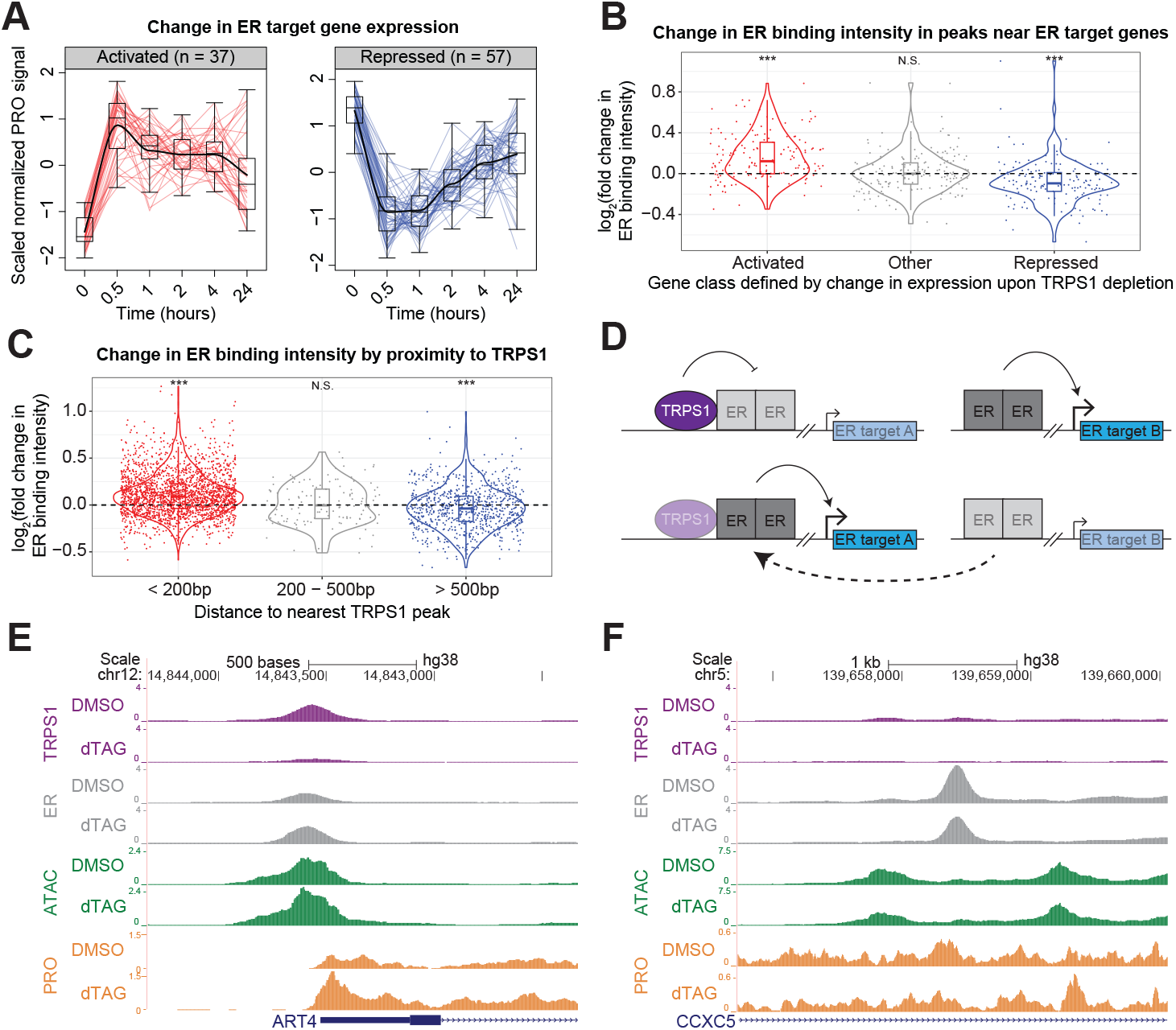
TRPS1 redistributes ER binding to modulate ER target gene transcription. A) Kinetic traces of activated and repressed ER target genes over the time course. B) Violin and box and whisker plots for ER binding intensity fold change upon TRPS1 depletion at ER ChIP-seq peaks within 100kb of the TSS of each gene, grouped by gene class defined by change in expression upon TRPS1 depletion. (In (B) and (C), *** represents a significant one-sample t-test p-value < 10^−3^. N.S. represents a non-significant p-value > 0.1.) C) Violin and box and whisker plots for ER binding intensity fold change upon TRPS1 depletion at ER ChIP-seq peaks, grouped by summit-to-summit distance to the nearest TRPS1 ChIP-seq peak. D) Model of TRPS1-mediated ER redistribution and modulation of ER target gene transcription. Transparent boxes indicate reduced binding intensity or attenuated transcription. Above, at baseline, TRPS1 directly decreases ER binding intensity proximal to TRPS1, attenuating ER activation of proximal ER target genes. Distal to TRPS1, ER binding intensity is not directly affected by TRPS1, and ER fully activates proximal ER target genes. Below, after TRPS1 depletion, ER binding proximal to TRPS1 increases in intensity, augmenting ER activation of proximal ER target genes. Distal to TRPS1, ER binding intensity is indirectly decreased, as limiting ER molecules are redistributed to TRPS1-proximal regulatory elements, attenuating ER activation of proximal ER target genes. E) ChIP, ATAC, and PRO density around an example increased ER binding site near an activated ER target gene. At this TRPS1-proximal ER binding site, upon dTAG treatment, TRPS1 binding intensity decreases, ER binding intensity increases, chromatin accessibility increases, and gene expression increases. F) ChIP, ATAC, and PRO density around an example decreased ER binding site near an repressed ER target gene. At this TRPS1-distal ER binding site, upon dTAG treatment, ER binding intensity decreases, chromatin accessibility decreases, and gene expression decreases. In (F) and (G), dTAG refers to dTAG-13 and dTAG^V^-1 at 50nM each.

We hypothesized that these changes in ER target gene transcription are mediated by changes in ER activity. We performed ER ChIP-seq to test the prediction that ER binding intensity proximal to dynamic ER target genes would change in concordance with the change in gene transcription. As we expected, we found that ER ChIP-seq peaks within a 100 kilobase (kb) window around the TSS of activated genes tended to increase in intensity, and ER ChIP-seq peaks within a 100kb window around the TSS of repressed genes tended to decrease in intensity (Figure 5B).

We further hypothesized that only the increased ER binding sites represent a direct effect of TRPS1 activity. Consistent with a prediction of this hypothesis, we found that ER binding proximal to TRPS1 tends to increase in intensity, and ER binding distal to TRPS1 tends to decrease in intensity (Figure 5C). Together, these data suggest a model in which TRPS1 depletion redistributes ER binding from TRPS1-distal sites to TRPS1-proximal sites and modulates ER target gene transcription proximal to the dynamic ER binding sites (Figure 5D). To illustrate this phenomenon, we provide an example of a TRPS1-proximal, increased ER peak near an activated ER target gene in (Figure 5E), and a TRPS1-distal, decreased ER peak near a repressed ER target gene in (Figure 5F).

### TRPS1 activity is associated with breast cancer patient outcomes

We next sought to connect the primary TRPS1-responsive genes with downstream cellular and patient-related outcomes. We defined a new steady state of transcription with the 24 hour time point after TRPS1 depletion. We ranked genes based on their shrunken fold change in PRO signal (Figure 6A). Using this ranking, we performed gene set enrichment analysis with the Hallmark gene sets and found multiple cell-cycle-related gene sets to be negatively enriched, including E2F Targets (Figure 6B) (Mootha et al. 2003; Subramanian et al. 2005). Consistent with this, we observed a significant decrease in cell number doubling rate of T47D dTAG-TRPS1 cells upon TRPS1 depletion (Figure 6C).

**Fig. 6.**
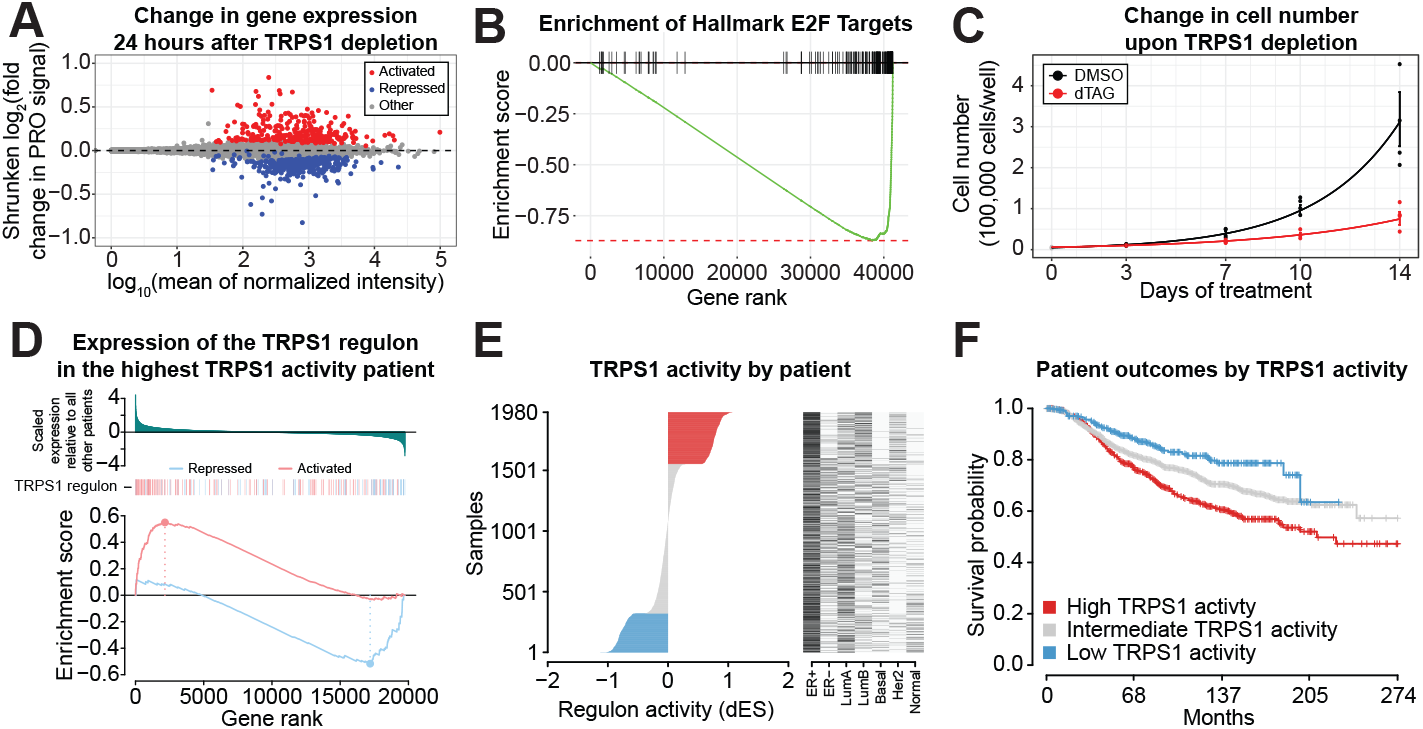
TRPS1 activity is associated with breast cancer patient outcomes. A) MA plot of PRO signal in each gene, with shrunken log fold change values representing transcription in the 30 minute dTAG-13 and dTAG^V^-1 at 50nM each (dTAG) treatment condition relative to the DMSO condition. B) Mountain plot of the Hallmark E2F Targets gene set, using genes ranked by shrunken fold change from (A). A negative enrichment score indicates an enrichment of the gene set among repressed genes. Adjusted p-value 2.5*10^−19^. C) Cell number over time of dTAG-TRPS1 cells treated with dTAG or DMSO. Analysis of variance for the coefficient corresponding to the difference in doubling rates between the conditions in a linear model of the logarithm of cell number versus time: p-value 1.1*10^−5^. D) Differential enrichment score (dES) calculation for an example patient with the highest TRPS1 activity. Above, genes ranked by scaled expression in this patient relative to all other patients in the METABRIC cohort (Curtis et al. 2012; Pereira et al. 2016). Below, gene set enrichment analysis of TRPS1-repressed and TRPS1-activated genes defined by response after 30 minutes of TRPS1 depletion. E) Patients from the METABRIC cohort, ranked by dES as calculated in (D), with classifications of the tumors on the right. F) Kaplan-Meier curves for patients in the METABRIC cohort, stratified by TRPS1 activity as in (E). Logrank p-value 8.3*10^−6^.

Finally, we calculated a TRPS1 activity score by adapting methods developed by (Castro et al. 2016; Fletcher et al. 2013). We used our PRO-seq data to determine a primary TRPS1 regulon based on the differentially expressed genes 30 minutes after TRPS1 depletion. We classified breast cancer patients from the METABRIC cohort as having high TRPS1 activity if both a) TRPS1-repressed genes are negatively enriched and b) TRPS1-activated genes are positively enriched, relative to all other patients in the cohort (example patient in (Figure 6D)) (Curtis et al. 2012; Pereira et al. 2016). Similarly, we classified patients as having low TRPS1 activity if both a) TRPS1-repressed genes are positively enriched and b) TRPS1-activated genes are negatively enriched. We classified the remaining patients as having intermediate TRPS1 activity. We ranked patients based on their TRPS1 activity and found no association with other clinical covariates (Figure 6E). When we stratified patients by TRPS1 activity, we found TRPS1 activity to be significantly associated with shorter survival time (Logrank p-value 8.3*10^−6^) (Figure 6F).

## Discussion

In this study, we used rapidly inducible targeted protein degradation to systematically determine the primary effects of acute TRPS1 depletion on chromatin accessibility, ER binding, and nascent transcription in a luminal breast cancer cell line. We focused on TRPS1 based on two orthogonal, genome-wide, unbiased assays that implicated TRPS1 in the processes of breast tumor incidence and breast cancer cell number accumulation.

First, we used the summary statistics from a recent GWAS to plot two sets of common genetic variants in the TRPS1 locus associated with breast cancer incidence (Zhang et al. 2020). These genetic variants were independently identified as significantly associated with breast cancer incidence in a previous GWAS (Michailidou et al. 2017), but Zhang *et al*. determined that the association was strongest among luminal breast tumors. Second, we analyzed data from the Cancer Dependency Map project and found that sensitivity to TRPS1 knockout was correlated with sensitivity to ER knockout and significantly enriched among luminal breast cancer cell lines (Meyers et al. 2017). Both of these unbiased screens indicate that TRPS1 contributes to luminal breast cancer cell fitness and led us to the hypothesis that TRPS1 influences ER activity.

As TFs regulate the transcription of many other chromatin-associated factors that themselves regulate RE activity, TF binding, and transcription, we sought to isolate the primary effects of TRPS1 depletion. To do so, we used the dTAG system to acutely deplete endogenous TRPS1 protein abundance within minutes of induction (Nabet et al. 2018). This is in contrast to traditional RNA interference or gene knockout methods, which can take days to deplete the target of interest.

Minutes to hours after TRPS1 depletion, we performed several sensitive, genome-wide assays. ATAC-seq and ChIP-seq can be performed at any time point after a perturbation, as they measure chromatin accessibility and chromatin-associated factor binding, which can change with rapid kinetics. In contrast, changes in messenger RNA abundance accumulate more slowly, with kinetics that depend not only on the rate of nascent transcription but also on the ratio of abundance to synthesis and degradation rates. In contrast, nascent transcriptional profiling measures the immediate change in RNA synthesis rates after a perturbation. Here we use PRO-seq coupled with acute TRPS1 depletion to identify primary TRPS1-responsive genes.

With our cell line and assays in hand, we first measured changes in chromatin accessibility upon TRPS1 depletion. Consistent with previous studies linking TRPS1 to corepressor complexes, we found that the predominant effect of TRPS1 depletion is an increase in chromatin accessibility and bidirectional transcription at REs (Cornelissen et al. 2020; Elster et al. 2018; Serandour et al. 2018; Wang et al. 2018a,b). We observed lower enrichment of the GATA motif prevalence in increased peaks over time, indicating that our shortest time point was the most specific for isolating the primary effects of TRPS1 depletion. Intriguingly, we identified a significant enrichment of ER half-site motifs in increased as well as decreased ATAC peaks, suggesting that ER activity was changing in a site-specific manner.

We next measured changes in nascent transcription minutes to hours after TRPS1 depletion and clustered the gene responses. Activated genes were enriched for cholesterol homeostasis genes. Of note, several recent GWAS have identified SNPs in the TRPS1 locus associated with blood cholesterol levels (Richardson et al. 2020; Ripatti et al. 2020; Sakaue et al. 2021). However, as of yet, no mechanistic follow-up studies into the role of TRPS1 in cholesterol biology have been performed. After TRPS1 depletion, repressed genes were enriched for estrogen response gene sets. Consistent with our previous data, activated genes were closer to increased ATAC peaks, suggesting TRPS1 directly represses these target genes at steady state. On the other hand, repressed genes were not closer to decreased ATAC peaks, suggesting a distinct mechanism of transcriptional regulation of this gene class.

We hypothesized that at steady state TRPS1 directly represses and indirectly activates its primary response genes. Using ER target genes and ER genomic binding as a case study, we found evidence supporting a model of acute ER redistribution. ER binding sites proximal to TRPS1 tended to increase in intensity upon TRPS1 knockdown, with distal ER binding sites tending to decrease in intensity. Furthermore, genes activated upon TRPS1 depletion were surrounded by ER binding sites that increased in intensity, and repressed genes were near decreased ER binding sites. Taken together, we propose a model in which TRPS1 directly decreases chromatin accessibility at steady state. Upon acute TRPS1 depletion, TRPS1-proximal REs increase in accessibility, allowing ER to redistribute from TRPS1-distal REs. Subsets of ER target genes are activated or repressed by TRPS1 depletion via distinct mechanisms.

First described in the 1980s, the concept of coactivator “squelching” has been debated as a mechanism of indirect activity distal from a TF’s genomic binding sites (Gill and Ptashne 1988; Tasset et al. 1990). Squelching has been proposed as a mechanism by which nuclear receptors like ER acutely repress transcription of a subset of primary response genes by competing for limiting coactivators (Bocquel et al. 1989; Guertin, MJ et al. 2014; Meyer et al. 1989; Schmidt, SF et al. 2016). Here we propose not the redistribution of coactivators by an activating TF, but a redistribution of activating TFs themselves via a rapid increase in local chromatin accessibility after the acute depletion of a repressive TF.

Our findings of both increased and decreased ER genomic binding and target gene transcription are distinct from previous studies of the effects of TRPS1 on TF binding and activity. Elster *et al*. used an unbiased screen to identify TRPS1 as a repressor of YAP activity in another luminal breast cancer cell line, MCF7 (Elster et al. 2018). After TRPS1 knockdown, the authors observed a genome-wide activation of YAP target genes. We did not find a YAP gene signature among dynamic genes in our PRO-seq data, though we did observe an enrichment of TEAD motifs in increased ATAC peaks, suggesting differences between the cells used in each study and perhaps their baseline YAP-TEAD activity. While we propose that our acute redistribution model is generalizable to other TFs and sets of TRPS1-regulated genes beyond ER and its target genes, it remains possible that TRPS1 modulates the activity of additional TFs in a unidirectional manner via a distinct mechanism.

Serandour *et al*. knocked down TRPS1 in MCF7 cells and reported both a genome-wide repression of ER target genes as well as a genome-wide increase in ER binding (Serandour et al. 2018). We did not perform ChIP-seq at a comparable time point to their days-long knockdown, so we cannot directly compare our ER binding data. Our latest PRO-seq time point was 24 hours after TRPS1 depletion, at which time we do not observe a genome-wide repression of ER target genes. This could once again be attributable to a difference in cell lines. However, we would also speculate that the unidirectional and nonconcordant changes in ER binding and target gene expression at later time points could be due to non-primary effects of extended TRPS1 knock-down.

Finally, we used PRO-seq data from both late and early time points to identify genes that represent cells at a new steady state after TRPS1 depletion, as well as primary TRPS1-responsive genes. After 24 hours of TRPS1 depletion, repressed genes were enriched for cell cycle related genes, consistent with a decrease in cell number doubling rate. Unique to this study, we used primary TRPS1-responsive genes to define a TRPS1 activity score, adapting a method based on predicted TF target genes (Castro et al. 2016; Fletcher et al. 2013). Using this method, we were able to stratify breast cancer patients into groups with differing survival probabilities.

Using TRPS1 activity score to classify patients may provide additional insight into the transcriptional program within a patient’s tumor that might not be immediately apparent based on previous surrogates for TRPS1 activity. For example, TRPS1 is frequently amplified in breast tumors, and this amplification is associated with worse prognosis (Radvanyi et al. 2005; Serandour et al. 2018). However, TRPS1 is often co-amplified along with the rest of the chromosomal segment 8q23–q24, where the proto-oncogene C-MYC resides, making it difficult to discern whether TRPS1 amplification is a driver of breast cancer progression (Savinainen et al. 2004).

In contrast, higher TRPS1 expression has been associated with better breast cancer patient outcomes, though its expression is highly correlated with ER and GATA3, both favorable prognostic indicators (Chen et al. 2010; Lin et al. 2017). As relative TF expression and activity across patients are not always identical, our data uses primary TRPS1-responsive genes as a measure of TRPS1 activity. Our TRPS1 activity score is not correlated with ER-positivity and effectively stratifies patients. Though our patient outcome analysis, as with all similar analyses, describes an association and does not necessarily imply a causative relationship, the direction is consistent with the effect on cell number observed in this study as well as the Cancer Dependency Map, suggesting that TRPS1 drives breast cancer cell number accumulation.

Altogether, we provide a systematic study of the primary effects of rapid TRPS1 depletion in luminal breast cancer cells. We propose a generalizable model in which TRPS1 depletion leads to decondensation of local chromatin structure, allowing for the acute redistribution of additional TFs like ER, both activating and repressing subsets of their target genes. This TRPS1-regulated transcription appears to be relevant for cancer cell fitness, as TRPS1 depletion decreases cell number doubling rate, and high TRPS1 activity is associated with worse breast cancer patient outcomes.

These methods of inducible targeted protein degradation coupled with genomic chromatin assays and nascent RNA transcriptional profiling should in principle be applicable to the study of any TF, allowing us to better understand the mechanisms behind the phenotypes associated with additional GWAS hits.

## Methods

### GWAS and DepMap data visualization

Summary statistics from (Zhang et al. 2020) were downloaded from the NHGRI-EBI GWAS Catalog (Sollis et al. 2023). SNPs in the TRPS1 locus were plotted using Locus-Zoom (Boughton et al. 2021). Knockout scores and luminal breast cancer identifiers were downloaded from the Cancer Dependency Map project (Meyers et al. 2017) and plotted using the statistical programming language R (R Core Team 2021).

### Cell culture

T47D cells (RRID:CVCL_0553) (ATCC) were cultured in RPMI 1640 medium (Gibco) supplemented with 10% fetal bovine serum (Gemini) and 10*µ*g/ml insulin from bovine pancreas (Sigma, made as a 1000x solution in 1% aqueous glacial acetic acid).

### Plasmid generation for gene editing

DNA for transfection was prepared as previously described (Sathyan et al. 2019, 2020). A CRISPR sgRNA (TTATCTTTGCAGATATGGTC) targeting the 5^*′*^ end of the TRPS1 coding sequences was designed using Benchling. The sgRNA was cloned into hSpCas9 plasmid PX458 (Addgene #48138) as previously described (Ran et al. 2013), using the following primers: 5^*′*^-CACCGTTATCTTTGCAGATATGGTC-3^*′*^ and 5^*′*^-AAACGACCATATCTGCAAAGATAAC-3^*′*^. A plasmid harboring a synthetic HygR-P2A-2xHA-FKBP_F36V insert was generated with Cold Fusion (System Biosciences), starting with the HygR-P2A-AID cassete in pMGS58 (Addgene #135311) (Sathyan et al. 2019) and the Puro-P2A-2xHA-FKBP_F36V casette in (Addgene #91793) (Nabet et al. 2018). The linear donor was generated by PCR using primers (IDT) that contain 50-nucleotide homology tails and gel-purified. The primers contained 5^*′*^ phosphorothioate modifications to increase PCR product stability in the cell (Zheng et al. 2014). The primers used for making PCR donor fragments were: 5^*′*^-G*T*AACTTTCAGATAACACTGTATCTGCCTTTT CCCTTTATCTTTGCAGATATGAAAAAGCCTGAACT CACCG-3^*′*^ and 5^*′*^-T*T*CACTTGCAACGTTTCTCAGAGGGGGGTTCT TTTTCCGGACACCTGAACCTGAACCTCCAGATCCACCAGATCTTTCCAGTTTTAGAAGCTCCACATCG-3^*′*^ with asterisks representing the phosphorothioate modifications.

### dTAG-TRPS1 clone generation

Clones were generated as previously described (Sathyan et al. 2019, 2020), with modifications. An initial round of cloning was performed using puromycin selection, but upon genomic DNA sequencing this clone did not appear to have a dTAG insertion event within TRPS1. Nevertheless, this clone was used for a second round of cloning using hygromycin selection. 3*10^6^ cells were plated in 10cm plates. The next day, cells were cotransfected with 15*µ*g of CRISPR/Cas9-sgRNA plasmid and 1.85*µ*g of linear donor PCR product using Lipofectamine 3000 (Thermo Fisher Scientific) in Optimem (Gibco). One day after transfection, the media was replaced. Starting four days after transfection, cells were selected for two weeks with 200*µ*g/mL of Hygromycin B (Invitrogen) with 20% conditioned media, replaced twice per week. Colonies were then grown in 20% conditioned media, replaced twice per week, until they were large enough to be picked and passaged to a 24-well plate. Clones were expanded and frozen at 8 passages after transfection. Integration was tested with Western blotting, PCR, and Sanger sequencing.

### Western blotting

8*10^5^ cells per sample were plated in each well of a 6-well plate. Cells were treated with DMSO or 50nM dTAG-13 and 50nM dTAG^V^-1 in DMSO at various time points and collected simultaneously. At the time of harvest, cells were scraped and lysed in RIPA buffer (1% Nonidet P-40, 1% sodium deoxycholate, 0.1% sodium dodecyl sulfate, 2mM EDTA, 150mM NaCl, 10mM sodium phosphate, 50mM NaF, 50mM Tris pH 7.5), with 100*µ*M benzamidine, 5*µ*g/mL aprotinin, 5*µ*g/mL leupeptin, 1*µ*g/mL pepstatin, 1mM phenylmethylsulfonyl fluoride, and 2mM sodium orthovanadate added fresh. Lysates were sonicated in a Biorupter UCD-200 (Diagenode) on high for 30 seconds on and 30 seconds off for 5 cycles, and clarified by centrifugation at 14,000rpm for 15 min in 4°C. Protein concentration was measured by BCA assay and diluted to the same concentration. 10× Laemmli buffer was added to a final concentration of 1x, and *β*-mercaptoethanol was added to a final concentration of 1%. Samples were boiled at 95°C for 10 minutes, and 30*µ*g of each was loaded into a 10% polyacrylamide gel. Samples were separated by gel electrophoresis and transferred to nitrocellulose membranes. Membranes were incubated in blocking buffer (3% bovine serum albumin, 1X Tris buffered saline) for 1 hour at room temperature with rocking. Primary antibodies (anti-TRPS1, Cell Signaling #17936S, and anti-*β*-Actin, Cell Signaling #3700S) were diluted 1:1,000 in primary buffer (3% bovine serum albumin, 0.1% sodium azide, 0.1% Tween-20, 1X Tris buffered saline) at 4°C with rocking overnight. Fluorescent secondary antibodies were diluted 1:10,000 in secondary buffer (5% bovine serum albumin, 0.1% sodium azide, 0.1% Tween-20, 1X Tris buffered saline) and incubated for 1 hour at room temperature with rocking, and fluorescence was measured (Odyssey, Licor).

### ChIP-seq library preparation

2.4*10^7^ cells per sample were plated across 3 15cm dishes 2 days before harvest. Cells were treated with DMSO or 50nM dTAG-13 and 50nM dTAG^V^-1 in DMSO for 30 minutes and collected simultaneously. At the time of harvest, cells were fixed with 1% formaldehyde (Sigma) for 10 minutes at 37°C and quenched with 125mM Glycine (Fisher) for 10 minutes at 37°C. Plates were moved to ice, and cells were washed and scraped into ice cold PBS containing Complete EDTA-free Protease Inhibitor Cocktail (Roche). Cells were pelleted in aliquots of 3.6*10^7^ cells, snap frozen in liquid nitrogen, and stored at -80°C. Pellets were thawed, and cells were lysed in 1mL Cell Lysis Buffer (85mM KCl, 0.5%NP40, 5mM PIPES pH 8.0), with protease inhibitor cocktail added fresh, for 10 minutes with rotation at 4°C. Nuclei were pelleted at 3300g at 4°C for 5 minutes and resuspended in 500*µ*L ChIP lysis buffer (0.5% SDS, 10mM EDTA, 50mM Tris-HCl pH 8.1), with protease inhibitor cocktail added fresh, for 10 minutes with rotation at 4°C. Lysates were moved to 15ml polystyrene conical tubes (Falcon) and sonicated in a Biorupter UCD-200 (Diagenode) on high for 30 seconds on and 30 seconds off for 4 sets of 5 cycles. Before each set, ice in the water bath was replaced, and samples were gently vortexed to mix. Sonicated lysates were then move to 1.5ml tubes and clarified by centrifugation at 14,000rpm for 15 min in 4°C. 500*µ*L of the supernatant was diluted into 6.5mL Dilution Buffer (0.01% SDS, 1.1% Triton X-100, 1.2mM EDTA, 167mM NaCl, 16.6mM Tris-HCl pH 8.0), with protease inhibitor cocktail added fresh (1*10^6^ cells in 200*µ*L). 1ml (5*10^6^ cells) was aliquoted into each of 3 tubes with antibody (1.25 *µ*g anti-HA, Cell Signaling #3724S, 2.5*µ*g anti-ER, Millipore #06-935, or 2.5*µ*g IgG control, Cell Signaling #2729S), and incubated with end-over-end rotation at 4°C overnight.

50*µ*L Protein A/G Magnetic Beads (Pierce) per sample were washed with bead washing buffer (PBS with 0.1% BSA and 2mM EDTA) and then incubated with samples for 2 hours with rotation at 4°C. The samples were washed once each with low salt immune complex buffer (0.1% SDS, 1% Triton x-100, 2mM EDTA, 150mM NaCl, 20mM Tris HCl pH 8.0), high salt immune complex buffer (0.1% SDS, 1% Triton x-100, 2mM EDTA, 500mM NaCl, 20mM Tris Hcl pH8.0), LiCl immune complex buffer (0.25M LiCl, 1% NP-40, 1% deoxycholate, 1mM EDTA, 10mM Tris-HCl pH8.0), and 1xTE (10mM Tris-HCl, 1mM EDTA pH8.0). Immune complexes were eluted in elution solution, (1% SDS, 0.1M sodium bicarbonate) in a thermomixer for 30 min at 65°C at 1,200rpm. Crosslinks were reversed and proteins were digested with the addition of 200mM NaCl and 2ul Proteinase K in a thermocycler at 65°C for 16 hours. DNA was purified with a Qiaquick PCR cleanup (Qiagen), and libraries were prepared with a NEBNext Ultra II Library Prep Kit (New England Biolabs).

### ChIP-seq analysis

Adapters were removed using `cutadapt` (Martin 2011). Reads were aligned to the *hg38* genome assembly with `bowtie2` (Langmead and Salzberg 2012). Duplicate reads were removed, and the remaining reads were sorted into *BAM* files and converted to *bed* format for counting with `samtools` (Li et al. 2009). Reads were also converted to *bigWig* format with `deeptools` (Ramírez et al. 2014).

Peaks were called with `MACS2` (Zhang et al. 2008). Reads were counted in peaks using `bedtools`, and differentially bound peaks were identified with `DESeq2` (Love et al. 2014; Quinlan and Hall 2010). Heatmaps were generated with `deeptools`. Peak proximity to and overlap with other features were calculated with `bedtools`.

### ATAC-seq library preparation

ATAC-seq libraries were prepared as previously described (Grandi et al. 2022), with modifications. 4 replicates were performed from cells treated and collected at different times in the same day. 4*10^5^ cells per sample were plated in each well of a 6-well plate 2 days before harvest. Cells were treated with DMSO or 50nM dTAG-13 and 50nM dTAG^V^-1 in DMSO at various time points and collected simultaneously. At the time of harvest, cells were moved to ice and scraped in 1mL ice cold PBS, and 100*µ*L (∼ 5 × 10^4^ cells) were transferred to 1.5 mL tubes. Cells were centrifuged at 500 x *g* for 5 minutes at 4°C, and the pellets were resuspended in 50 *µ*L cold lysis buffer (10mM Tris-HCl, 10mM NaCl, 3mM MgCl_2_, 0.1% NP-40, 0.1% Tween-20, 0.01% Digitonin, adjusted to pH 7.4) and incubated on ice for 3 minutes. Samples were washed with 1 mL cold wash buffer (10mM Tris-HCl, 10mM NaCl, 3mM MgCl_2_, 0.1% Tween-20). Cells were centrifuged at 500 x *g* for 10 minutes at 4°C, and pellets were resuspended in the transposition reaction mix (25 *µ*L 2X TD buffer (Illumina), 2.5 *µ*L TDE1 Tn5 transposase (Illumina), 16.5 *µ*L PBS, 0.5 *µ*L 1% Digitonin, 0.5 *µ*L 10% Tween-20, 5 *µ*L nuclease-free water) and incubated in a thermomixer at 37°C and 100rpm for 30 minutes. DNA was extracted with the DNA Clean and Concentrator-5 Kit (Zymo Research). Sequencing adapters were attached to the transposed DNA fragments using NEBNext Ultra II Q5 PCR mix (New England Biolabs), and libraries were amplified with 8 cycles of PCR. PEG-mediated size fractionation (Lis 1980) was performed on the libraries by mixing SPRIselect beads (Beckman) with each sample at a 0.5:1 ratio, then placing the reaction vessels on a magnetic stand. The right side selected sample was transferred to a new reaction vessel, and more beads were added for a final ratio of 1.8:1. The final size-selected sample was eluted into nuclease-free water. This size selection protocol was repeated to further remove large fragments.

### ATAC-seq analysis

Adapters were removed using `cutadapt` (Martin 2011). Reads aligning to the mitochondrial genome with `bowtie2` (Langmead and Salzberg 2012) were removed. The remaining reads were aligned to the *hg38* genome assembly with `bowtie2`. Duplicate reads were removed, and the remaining reads were sorted into *BAM* files with `samtools` (Li et al. 2009). Reads were converted to *bed* format with `seqOutBias` and *bigWig* format with `deeptools` (Martins et al. 2018; Ramírez et al. 2014). Accessibility peaks were called with `MACS2` (Zhang et al. 2008). Reads were counted in peaks using `bedtools`, and differentially accessible peaks were identified with `DESeq2` (Love et al. 2014; Quinlan and Hall 2010). *de novo* motif identification was performed on dynamic peaks with `MEME`, and `TOMTOM` was used to match motifs to the HOMER, Jaspar, and Uniprobe TF binding motif databases (Bailey et al. 2015; Heinz et al. 2010; Hume et al. 2015; Khan et al. 2018).

AME was used to identify motifs enriched in increased or decreased peaks relative to unchanged peaks (McLeay and Bailey 2010). `FIMO` and `bedtools` were used to assess motif enrichment around peak summits (Grant et al. 2011). Dynamic peaks were clustered into response groups using `DEGreport` (Pantano 2019).

### PRO-seq library preparation

Cell permeabilization was performed as previously described (Mahat et al. 2016), with modifications. 4 replicates were performed from cells treated and collected at different times in the same day. 8*10^6^ cells per sample were plated in 15cm dishes 2 days before harvest. Cells were treated with DMSO or 50nM dTAG-13 and 50nM dTAG^V^-1 in DMSO at various time points and collected simultaneously. At the time of harvest, cells were scraped in 10mL ice cold PBS and washed in 5mL buffer W (10mM Tris-HCl pH 7.5, 10mM KCl, 150mM sucrose, 5mM MgCl_2_, 0.5mM CaCl_2_, 0.5mM DTT, 0.004U/mL SUPERaseIN RNase inhibitor (Invitrogen), Complete protease inhibitors (Roche)). Cells were permeabilized by incubating with buffer P (10 mM Tris-HCl pH 7.5, KCl 10 mM, 250 mM sucrose, 5 mM MgCl_2_, 1 mM EGTA, 0.05% Tween-20, 0.1% NP40, 0.5 mM DTT, 0.004 units/mL SUPERaseIN RNase inhibitor (Invitrogen), Complete protease inhibitors (Roche)) for 3 minutes on ice. Cells were washed with 10 mL buffer W before being transferred into 1.5mL tubes using wide bore pipette tips. Finally, cells were resuspended in 50*µ*L buffer F (50mM Tris-HCl pH 8, 5mM MgCl_2_, 0.1mM EDTA, 50% Glycerol, 0.5 mM DTT). Cells were snap frozen in liquid nitrogen and stored at -80°C.

PRO-seq libraries were prepared as previously described (Judd et al. 2020), with modifications. RNA extraction after the run-on reaction was performed with 500*µ*L Trizol LS (Thermo Fisher) followed by 130*µ*L chloroform (Sigma). The equivalent of 1*µ*L of 50*µ*M for each adapter was used. A random eight base unique molecular identifier (UMI) was included at the 5^*′*^ end of the adapter ligated to the 3^*′*^ end of the nascent RNA. 37°C incubations were performed with rotation with 1.5mL tubes placed in 50mL conical tubes in a hybridization oven. For the reverse transcription reaction, RP1 was used at 100*µ*M and dNTP mix was used at 10mM each. Libraries were amplified by PCR for a total of 8 cycles in 100*µ*L reactions with Phusion polymerase (New England Biolabs). No PAGE purification was performed to ensure that our libraries were not biased against short nascent RNA insertions.

### PRO-seq analysis

Adapters were removed using `cutadapt` (Martin 2011). Libraries were deduplicated using `fqdedup` and the 3^*′*^ UMIs (Martins and Guertin 2018). UMIs were removed, and reads were reverse complemented with the `seqtk`. Reads aligning to the rDNA genome with `bowtie2` (Lang-mead and Salzberg 2012) were removed. The remaining reads were aligned, sorted, and convert to *bed* and *bigWig* files with `bowtie2`, `samtools`, `seqOutBias`, and `deeptools`, respectively (Li et al. 2009; Martins et al. 2018; Ramírez et al. 2014). Composite profiles around TRPS1 peaks were generated with `deeptools`. Reads were counted in genes using `bedtools`, and differentially expressed genes were identified with `DESeq2` (Love et al. 2014; Quinlan and Hall 2010). Dynamic genes were clustered into response groups using `DEGreport` (Pantano 2019). Over-representation analysis was performed with `enrichr` (Chen et al. 2013), and gene set enrichment analysis was performed with `fgsea` (Korotkevich et al. 2021), both using the Hallmark gene sets (Liberzon et al. 2015).

### Genome browser visualization

Genome browser (Kent et al. 2002) images were taken from the following track hub: https://data.cyverse.org/dav-anon/iplant/home/TBD

### Cell number enumeration

1.25*10^4^ cells per sample were plated in a 24-well plate. The next day (day 0), cells were treated with DMSO or 50nM dTAG-13 and 50nM dTAG^V^-1 in DMSO. Media was replaced, maintaining the treatment condition, every 2 days. Cells were enumerated using a hemocytometer. 2 technical replicates were used on day 0, 2 for each treatment on day 3, and 3 for each treatment on days 7, 10, 14. The technical replicates were merged, and the experiment was performed in 4 biological replicates from different cell passages. The data were imported into R (R Core Team 2021) for visualization and statistical analysis. A linear model was fit for the log-transformed cell number and the time. A second linear model was fit that included an interaction term between the time and the treatment condition, representing the effect of treatment on the doubling rate. Analysis of variance was performed on the two models to test for the significance of the interaction term.

### TRPS1 activity score and patient outcome stratification

Primary TRPS1-regulated genes were defined based on the 30 minute time point using `DESeq2` (Love et al. 2014).

This TRPS1 regulon was then used in `RTN` (Castro et al. 2016; Fletcher et al. 2013) to define a TRPS1 activity score for each patient within the METABRIC cohort (Curtis et al. 2012; Pereira et al. 2016).

## Data Access

All analysis details and code are available at https://guertinlab.github.io/TRPS1_ER_analysis/Vignette.html. Raw sequencing files and processed counts and *bigWig* files are available from GEO SuperSeries accession record GSE236176, with SubSeries accession records GSE236175 (ATAC-seq), GSE236174 (ChIP-seq), and GSE236172 (PRO-seq).

## ACKNOWLEDGEMENTS

This work was funded by R35-GM128635 to MJG. 5T32GM007267-39 and 5T32GM008136-35 supported TGS. We thank Arun Dutta, Jacob Wolpe, Devin Roller, and Adam Spencer for critical feedback.

## AUTHOR CONTRIBUTIONS

All authors contributed to the conceptualization of the project. SKM contributed to the methodology. TGS performed the experiments, analyzed the data, and wrote the original draft of the manuscript. All authors reviewed and edited the manuscript.

---

**Fig. S1.**
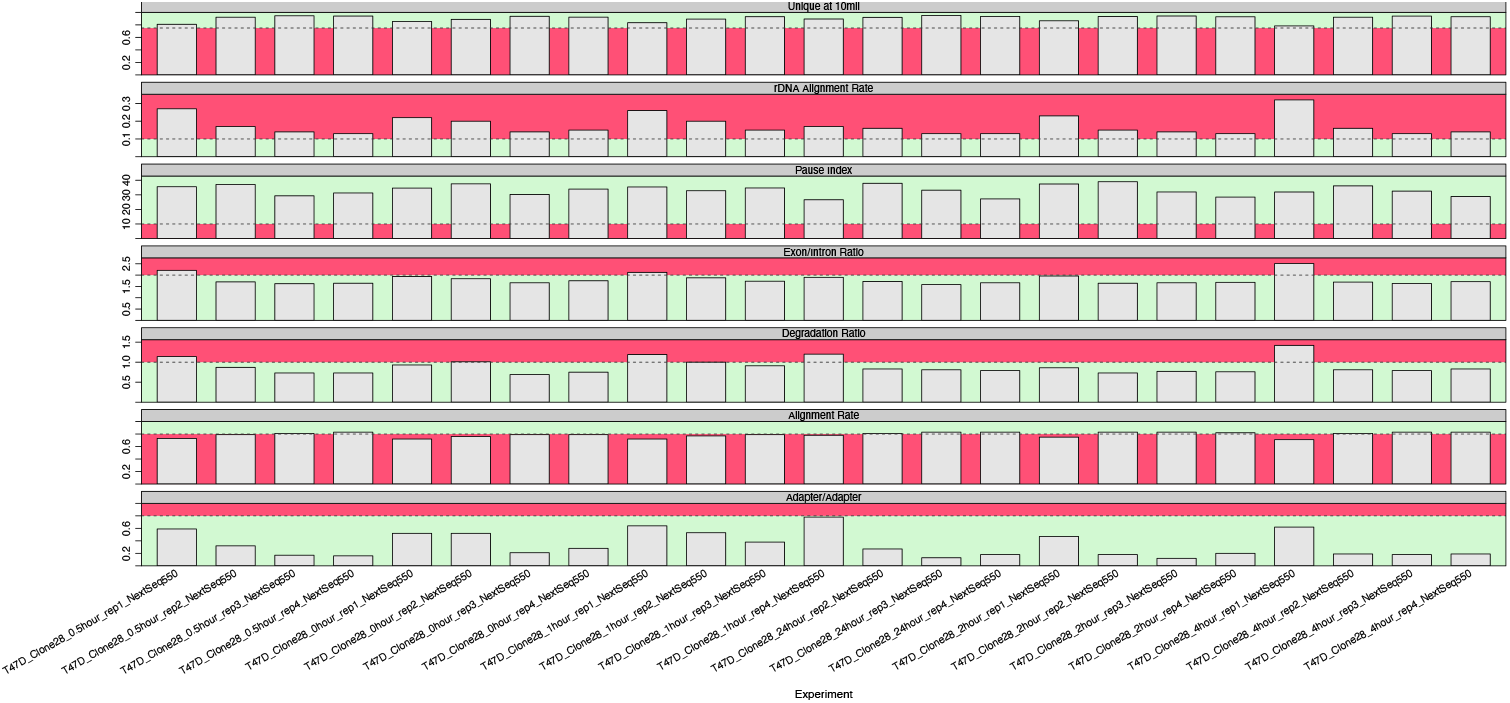
Quality control metrics for PRO-seq libraries. Quality control metrics are defined as in (Scott et al. 2022). Each metric is a row, and each sample is a column. The green region for each metric is the goal for a high quality library.

**Fig. S2.**
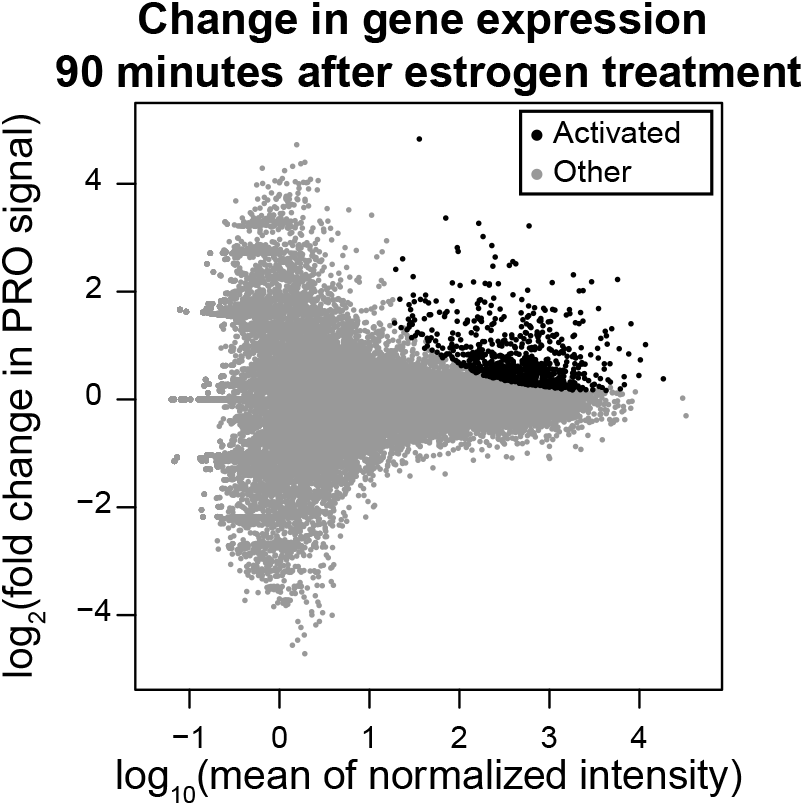
Acute estrogen treatment identifies direct ER target genes in T47D cells. MA plot of PRO signal, with fold change values representing transcription in the 90 minute estrogen treatment condition relative to the DMSO condition. Each point represents a gene, and black points represent the estrogen-activated genes that we use in Figure 5.

**Table S1.** Top 50 motifs significantly enriched in increased ATAC peaks relative to unchanged ATAC peaks at 30 minutes. Results generated using AME (McLeay and Bailey 2010).

| Motif | Consensus | Percent increased | Percent unchanged | Padj |
| --- | --- | --- | --- | --- |
| GATA5-Jaspar | WGATAASR | 69.70 | 8.47 | 2.95e-88 |
| GATA3-Jaspar | WGATAASR | 62.29 | 6.99 | 6.34e-76 |
| GATA3-Homer | AGATAASR | 66.10 | 9.96 | 8.55e-74 |
| TRPS1-Homer | AGATAAVANN | 61.86 | 8.05 | 6.96e-71 |
| GATA2-Jaspar | NBCTTATCTNH | 54.45 | 5.30 | 1.76e-65 |
| Gata6-Homer | BCTTATCWNN | 56.78 | 6.99 | 9.67e-64 |
| Gata4-Homer | NBWGATAAGV | 54.24 | 6.57 | 4.83e-60 |
| GATA6-Jaspar | NDNAGATAAGADD | 48.52 | 5.08 | 2.20e-54 |
| Gata2-Homer | BBCTTATCTS | 46.19 | 4.24 | 2.54e-53 |
| Gata6-Secondary-Uniprobe | WHNVDWGATAAGADTHN | 45.13 | 3.81 | 4.37e-53 |
| Gata1-Jaspar | TYCTTATCTSY | 41.95 | 3.18 | 7.08e-50 |
| Gata1-Homer | SAGATAAGVV | 38.56 | 3.18 | 9.24e-44 |
| Gata5-Secondary-Uniprobe | HWWRCTGATAAGRRNAN | 40.47 | 4.66 | 2.71e-41 |
| Gata4-Jaspar | YCTTATCTSHB | 40.47 | 5.08 | 9.06e-40 |
| Gata3-Secondary-Uniprobe | BDWDDAKAGATAAGARWTDARD | 31.99 | 2.75 | 2.39e-34 |
| TEAD3-Jaspar | RCATTCCW | 39.83 | 11.02 | 7.01e-23 |
| Mtf1-Secondary-Uniprobe | ANATAWGAAAWADV | 49.79 | 18.86 | 1.19e-21 |
| FOXP3-Jaspar | RTAAACA | 48.94 | 18.22 | 1.31e-21 |
| Arid3a-Secondary-Uniprobe | NNVVRTATCRDRWHD | 29.87 | 5.93 | 5.62e-21 |
| FO XK1-Homer | NNNTGTTTAY | 47.46 | 17.58 | 9.71e-21 |
| TEAD4-Jaspar | NRCATTCCWN | 41.53 | 13.77 | 9.84e-20 |
| HLTF-Jaspar | NHMCWTDKNN | 52.54 | 22.46 | 1.75e-19 |
| TEAD1-Jaspar | YRCATTCCWH | 40.04 | 13.35 | 1.34e-18 |
| FOXD2-Jaspar | GTAAACA | 49.15 | 20.55 | 3.63e-18 |
| FOXB1-Jaspar | WATGTAAATAT | 31.78 | 9.11 | 2.21e-16 |
| TEAD4-Homer | SSWGGATGY | 47.67 | 20.76 | 4.13e-16 |
| Glis2-Secondary-Uniprobe | HDTATTAWTAAAGV | 23.73 | 4.87 | 1.57e-15 |
| FOXM1-Homer | TRTTTRCYYW | 47.03 | 20.97 | 4.30e-15 |
| Arid3a-Jaspar | ATYAAA | 30.08 | 8.90 | 8.85e-15 |
| FOXL1-Jaspar | RTAAACA | 42.16 | 17.37 | 9.78e-15 |
| Prop1-Homer | NTAATBNVATTA | 27.97 | 7.63 | 9.94e-15 |
| FOXC1-Jaspar | WAWGTAAAYAW | 34.53 | 12.08 | 2.36e-14 |
| TEAD2-Jaspar | NYACATTCCWNS | 38.56 | 15.25 | 7.03e-14 |
| Sox5-Secondary-Uniprobe | NHDCATMATTDASNN | 27.33 | 7.84 | 1.43e-13 |
| FoxD3-Homer | TGTTTAYTTWGC | 36.65 | 14.19 | 2.42e-13 |
| Foxj2-Jaspar | RTAAACAA | 31.57 | 10.81 | 3.93e-13 |
| TEAD3-Homer | YRCATTCCAG | 27.54 | 8.26 | 5.35e-13 |
| Hoxa11-Homer | TTTTATDRCH | 29.03 | 9.32 | 8.93e-13 |
| NR2F2-Jaspar | NAAAGGTCANR | 24.79 | 6.78 | 1.40e-12 |
| FOXC2-Jaspar | WAWGTAAACAWW | 36.65 | 14.83 | 1.88e-12 |
| Zfp128-Secondary-Uniprobe | NGTATAYDTATAMN | 20.55 | 4.66 | 4.07e-12 |
| Barx1-Homer | AAACMATTAN | 26.48 | 8.47 | 1.80e-11 |
| FOXA1-Homer | WAWGTAAACA | 31.99 | 12.29 | 2.98e-11 |
| Hoxd11-Homer | NSYMATAAAA | 30.08 | 11.23 | 6.99e-11 |
| TEAD2-Homer | SCWGGATGY | 29.45 | 10.81 | 7.23e-11 |
| Fox:Ebox-Homer | NNNVCWGWGYAAACAVN | 36.44 | 15.89 | 8.97e-11 |
| GSC-Homer | RGGATTAR | 24.79 | 7.84 | 9.75e-11 |
| Pitx1-Jaspar | YTAATCCH | 24.36 | 7.63 | 1.19e-10 |
| FOXO4-Jaspar | GTAAACA | 33.05 | 13.56 | 1.61e-10 |

**Table S2.**
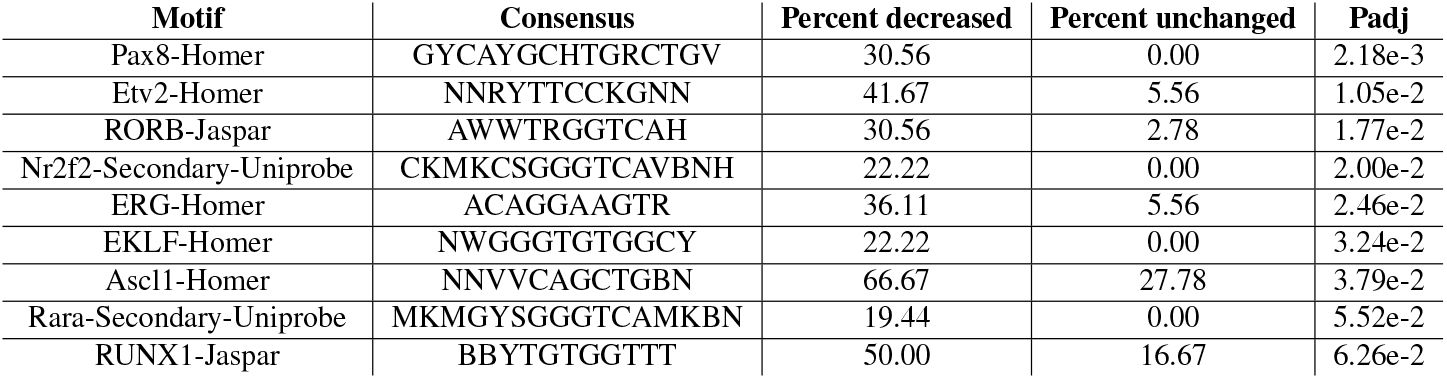
Motifs significantly enriched in decreased ATAC peaks relative to unchanged ATAC peaks at 30 minutes. Results generated using AME (McLeay and Bailey 2010).

